# Investigating the Mechanisms and Potential Therapeutic Targets of Vestibular Migraine

**DOI:** 10.1101/2025.01.28.635232

**Authors:** Xiaotong Yao, Zhuoji Liang, Yanling Liang

**Affiliations:** Department of Neurology; Guangdong Provincial Key Laboratory of Major Obstetric Diseases; Guangdong Provincial Clinical Research Center for Obstetrics and Gynecology; The Third Affiliated Hospital, Guangzhou Medical University, Guangzhou, 510150, Guangdong, China

**Keywords:** vestibular migraine,pathophysiology, feature genes, bioinformatics, therapeutic targets

## Abstract

**Background:** Vestibular Migraine (VM) is a complex neurological disorder with recurrent headaches and various vestibular symptoms. Although it severely affects patients’ quality of life, its causes and pathophysiology are still unclear, and effective treatments are scarce. The lack of data emphasizes the need for bioinformatics to find key genes and pathways in VM, which could help develop new diagnostic and treatment methods.

**Method:** The GSE109558 dataset was acquired from the Gene Expression Omnibus (GEO) database. To identify VM - related differentially expressed genes, screening was carried out through limma and Weighted Correlation Network Analysis (WGCNA).The functional analysis of VM - related differentially expressed genes was conducted using three bioinformatics approaches: Gene Ontology (GO) analysis, Kyoto Encyclopedia of Genes and Genomes (KEGG) pathway analysis, and Gene Set Enrichment Analysis (GSEA). Feature selection was further refined using lasso regression and random forest. Also, CIBERSORT was utilized to analyze the infiltration of immune cells, and Spearman’s correlation analysis was employed to explore the correlations between diagnostic differentially expressed genes and immune cells.Finally, the Comparative Toxicogenomics Database (CTD) was utilized to search for corresponding drugs, and molecular docking was performed to explore potential therapeutic targets.

**Result:** Six key feature genes (CABIN1, IFIT3, HEATR1, ARHGDIA, RAB11FIP4, and ZNF444) were identified as potential diagnostic markers for VM. Among these, CABIN1 demonstrated the highest diagnostic potential based on ROC curve performance, highlighting its promise as a diagnostic biomarker.Functional annotation of DEGs revealed their enrichment in biological processes related to inflammation, calcium ion channel regulation, and other pathways likely involved in VM pathophysiology. Through the CTD, drugs like Acetaminophen, bisphenol A, and Phenylephrine were identified.

Molecular docking simulation was used to explore their potential therapeutic mechanisms for VM.

**Conclusion:** This study offers important insights into the molecular mechanisms of VM and identifies six key feature genes, with CABIN1 standing out as a potential diagnostic marker.These findings pave the way for further research to validate the diagnostic and therapeutic implications of these genes and pathways.

## Introduction

Vestibular Migraine (VM) is the second most common cause of episodic vestibular syndrome, and patients often present to neurology with recurrent episodes of vertigo and dizziness as their main complaints, showing vestibular symptoms associated with headache, accompanied by nausea, vomiting, tinnitus, photophobia, etc[1]. Triggers of VM attacks often include stress, menstruation, irregular sleep, climate change, exertion, diet, etc[2]. Because of the heterogeneity of Vestibular Migraine , its manifestations vary greatly among individuals, so the rate of misdiagnosis and underdiagnosis is relatively high. Because of the heterogeneity of VM, the manifestation of Vestibular Migraine varies greatly among individuals, and it is often described as the “chameleon” of vestibular diseases, so the misdiagnosis and underdiagnosis rate are high[3]. It has been suggested that Vestibular Migraine has a tendency to become chronic, which may affect cognitive function and lead to mood disorders[4]. The underlying pathophysiological mechanisms of VM are complex and controversial, with several potential mechanisms proposed, including ion channel dysfunction,central sensitization, and neuroinflammation.

Given the absence of effective treatments, uncovering efficacious therapeutic targets is of utmost importance. A growing body of evidence indicates that inflammation, manifested by glial cell activation and leukocyte infiltration, has a pivotal role in the development of Vestibular Migraine[5, 6]. In addition, inflammatory processes are also believed to play a role in certain syndromes associated with Vestibular Migraine, such as anxiety and depression, as well as common behavioral manifestations (e.g., sleep disturbances, reduced activity, and decreased social interaction)[2]. There is evidence that inflammatory mediators, including pro-inflammatory cytokines and chemokines, play a crucial role in the development and maintenance of Vestibular Migraine[7] . Microarray represents a firmly established approach employed for high - throughput, genome - wide gene expression profiling. In the realm of modern biology, bioinformatics offers a plethora of tools that are instrumental in analyzing the extensive volume of information housed within GEO databases. This analysis is crucial as it enables the identification of pivotal genes and critical pathways, which can, in turn, serve as the foundation for developing novel clinical therapeutic strategies.Against this backdrop, this study aims to deeply explore the molecular mechanisms and pathogenesis of VM and is committed to providing a new basis for the prevention and diagnosis of this complex disorder.

## Materials and Methods

### Microarray Datasets of Vestibular Migraine

Gene expression microarray GSE109558 (26 samples: 15 Vestibular Migraine (VM)s and 11 controls,) was downloaded from the National Center for Biotechnology Information (NCBI) subdataset gene expression Omnibus (GEO)database(http://www.ncbi.nlm.nih.gov/geo, accessed November 5, 2023) accessed on November 5, 2023) was downloaded, which includes bioexpression data for many species obtained through arrays, SNP arrays, and high-throughput sequencing .

The datasets used in this study were all retrieved from public databases. In order to guarantee the integrity and dependability of our analysis, a comprehensive inspection of the raw gene expression data from the GEO dataset was carried out by means of box plots. Following the removal of outliers, our final analytical dataset consisted of 25 samples(15 individuals with Vestibular Migraine and 10 controls).These public databases allow researchers to analyze for scientific purposes and therefore do not require ethical approval.

### Differential expression analysis

The gene expression matrix, platform information, and clinical data were retrieved from the GEO database. Probe IDs in the expression matrix were then mapped to gene symbols using the corresponding platform data, resulting in a globally standardized expression matrix with gene symbols as row identifiers and sample names as column identifiers[8]. If multiple probe sets correspond to the same gene, their average values are calculated.

Next, we further analysed the DEGs in the Vestibular Migraine and control groups using the R language limma package. The results are presented in the form of an image, listing the top 118 genes sorted by p - value. The image is derived from the results of the Wald test performed when comparing the samples of the two groups. After screening based on the default adjusted p - value threshold of 0.05, genes with significantly differential expression are highlighted. Specifically, up - regulated genes are marked in red, while down - regulated genes are shown in green. DEGs refer to those genes with a p - value < 0.05 and a fold - change (FC) ≥ 1.In addition, we utilized the weighted gene co-expression network analysis to uncover gene regulatory mechanisms and functional modules. During the data preprocessing stage, outliers and abnormal samples were removed. The correlation coefficients between genes were calculated to construct a correlation matrix. The scale-free network topology modeling function of the WGCNA package was employed to select an appropriate soft-thresholding power, transforming the correlation matrix into a weighted adjacency matrix and calculating the topological overlap matrix (TOM). Based on the TOM matrix, hierarchical clustering analysis was performed using dissimilarity as a similarity measure to identify key gene modules associated with VM.

### Gene Ontology and Pathway Enrichment Analysis

DEGs identified within the significant modules were subjected to gene enrichment analysis using the ClusterProfiler package in R (version 4.3.1). The analysis encompassed GO and KEGG pathway enrichment to elucidate the biological functions and molecular pathways of the DEGs. The enrichment results were subsequently visualized through the enrichplot package to facilitate a comprehensive interpretation of the underlying biological processes.

### Gene set enrichment analysis

GSEA uses pre-defined gene sets to rank genes based on their differential expression levels between two types of samples, and then checks whether the pre-set gene set is enriched at the top or bottom of this ranking table[9, 10]. The enrichment degree of GSEA was evaluated by using the normalized enrichment score (NES), and the potential enrichment pathways of vestibular migraine - related biological processes were deeply explored. The significance of enrichment was assessed under the conditions that the false discovery rate (FDR) was less than 0.25 and the p - value was less than 0.05.

### Identification of hub genes via machine learning

To further screen for hub genes in Vestibular Migraine, we employed two machine - learning algorithms: LASSO (Least Absolute Shrinkage and Selection Operator) and RF (Random Forest)Through the LASSO algorithm, we identified 6 potential biomarker candidates. The Random Forest algorithm then ranked these candidates. We selected genes based on the importance calculation of each gene and picked the top 30 as potential candidate genes for Vestibular Migraine. By comparing the results of different algorithms, we finally determined six key genes. The core genes of Vestibular Migraine include: CABIN1, IFIT3, HEATR1, ARHGDIA, RAB11FIP4, and ZNF444.

Subsequently, we plotted the Receiver Operating Characteristic (ROC) curves for these genes to evaluate the diagnostic efficacy of these core genes. The visualization results are presented as shown in the figure.

### Analysis of Immune Cell Composition in Vestibular Migraine Patients Using the CIBERSORT Algorithm

Utilizing the CIBERSORT algorithm, an RNA-seq based deconvolution method, this study analyzes the immune cell composition in patients with vestibular migraine compared to a healthy control group. CIBERSORT estimates the proportions of various immune cell subtypes within complex tissues[11]. By contrasting immune cell proportions between the VM and control groups, the study reveals distinctive immune cell features associated with VM. Additionally, correlation heatmaps are employed to further investigate the interrelationships among immune cell subtypes, aiming to identify potential immune regulatory mechanisms. The methodological advantage of CIBERSORT lies in its ability to precisely quantify the relative proportions of immune cell types, thereby facilitating a deeper understanding of the immunopathological mechanisms underlying the disease.

### Construction of transcription factor gene networks

The identified hub genes were imported into the NetworkAnalyst platform (version 3.0, https://www.networkanalyst.ca/) to construct transcription factor (TF)-gene interaction networks[12]. The purpose of this analysis was to uncover the regulatory relationships between transcription factors and target genes, thereby gaining deeper insights into the potential regulatory mechanisms of the hub genes. Furthermore, it aimed to identify key transcription factors that may influence the expression and functions of these genes, providing critical clues for elucidating their roles in specific biological processes or disease mechanisms.

### Identification of candidate drugs

For the key differentially expressed genes selected for vestibular migraine , we carried out predictions on potential drug molecules by leveraging the Comparative Toxicogenomics Database (CTD). We retrieved the protein structures of the key genes and those of the drugs from the RCSB Protein Data Bank (RCSB PDB, website: https://www.rcsb.org/) and the PubChem database (https://pubchem.ncbi.nlm.nih.gov/). The collected data were processed according to a previously published study and then presented in a visual form. Subsequently, we used Pymol software to determine the affinity parameters and three - dimensional spatial structures

## Results

### Identification of Vestibular Migraine-Associated Genes

The Metrix series, along with its corresponding platform information (GPL10558) and clinical data, were downloaded from the NCBI website. This dataset contains high-throughput gene expression data and microarray data, with the accession number GSE109558. Based on the study’s objectives, 25 samples were selected for in-depth analysis (15 from the VM group and 10 from the Control group).Differential analyses of genes in the GSE were performed in order to find genes closely associated with Vestibular Migraine. The deg of the dataset was found using the limma tool with a cut-off criterion of p < 0.05, |log (fold change)| ≥ 1. The results showed a total of 118 deg (26 up-regulated and 93 up-regulated). The details are shown in (Figure 1a,b). Based on the identified differentially expressed genes, we constructed a heatmap to show the expression patterns of these genes in the GEO dataset, with coloring based on the samples for easier analysis(Figure 1c).

**Figure 1.**
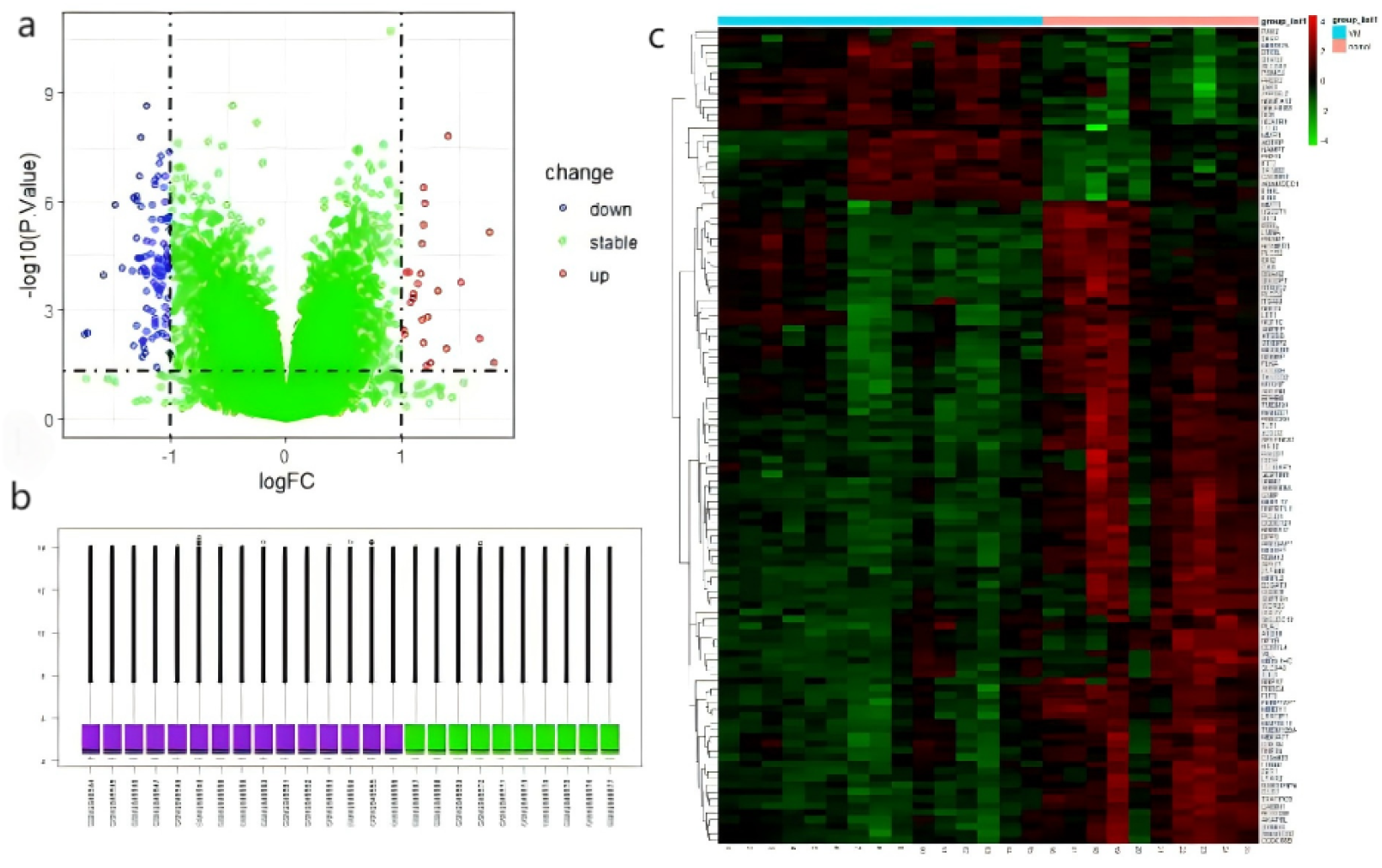
(A) deg identification between Vestibular Migraine (VM) and Control groups in the GSE 109558 dataset. Highlighted genes were significantly differentially expressed at the default p-value cut-off of 0.05 (red = up-regulated, green = down-regulated).(B) Box plot of raw gene expression in the GEO dataset.(C)Heatmap based on gene expression dataset. p < 0.05, |log(fold change)| ≥ 1.

### WGCNA and key module identification

Utilizing WGCNA, key gene modules in polyarticular VM were identified.As shown in Fig. 2A, when R^2^ > 0.85 and the average connectivity was high, the soft threshold was set to 24. Ten modules were identified using the dynamic tree-cut method (Figure 2B).

**Figure 2:**
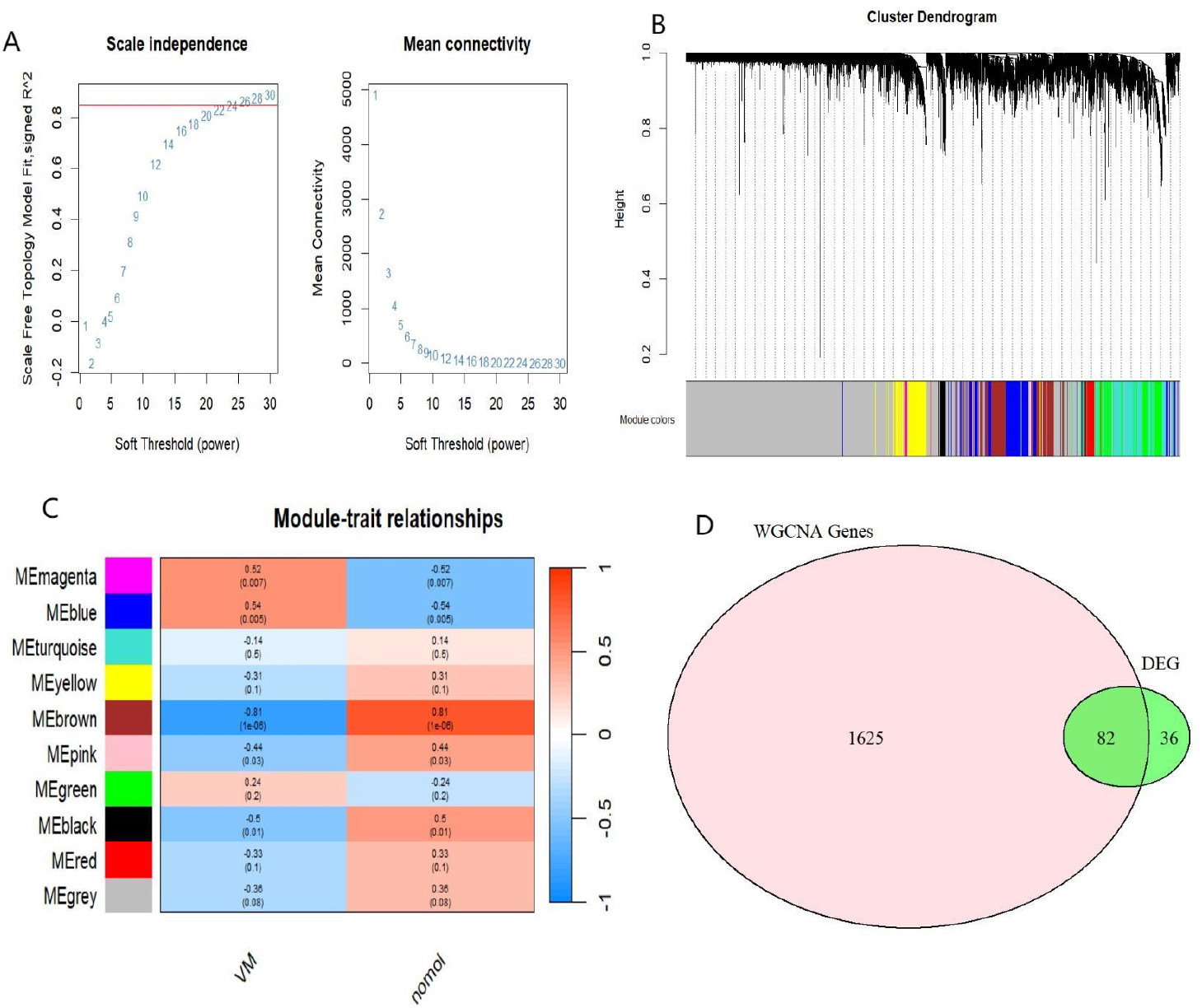
WGCNA analysis results. (A) Soft threshold and scale-free topology fitting index; (B) Combination module under cluster tree; (C) Module trait correlation heatmap. Red represents positive correlations, while blue represents negative correlations;(D)Venn Diagram and Intersection Display of DEGS Genes and WGCNA Genes.Pink represents WGCNA, and green represents DEGs.

By establishing module-trait relationships, we determined the correlation between modules and disease occurrence. As depicted in Fig. 2C, the blue module, which encompasses 882 genes, exhibited a significant positive correlation with septic shock (R = 0.54, p < 0.05). Conversely, the brown module, consisting of 825 genes, was significantly negatively associated with the disease (R = -0.81, p < 0.05).Based on this, through WGCNA, it was determined that both the blue module and the brown module were associated with VM. Finally, 82 genes were identified in the intersection of the screened core module genes and differentially expressed genes.

### Enrichment analysis

GO functional annotation and KEGG pathway enrichment analysis were meticulously conducted on the differentially expressed genes. For biological processes (bp), these DEGs were mainly involved in biological processes including neutrophil degranulation, neutrophil activation, involvement in immune responses, protein localisation to cell-cell junctions and neutrophil-mediated immunity. These processes are primarily associated with immune responses and cellular communication. Significant enrichment of cellular components includes transferrin transport, tertiary granules, phagocytic cups, tertiary granule membranes, endocytosed vesicles and endocytosed vesicle membranes. These components are associated with cellular structures involved in immune cell function and intracellular transport. Molecularfunctions include serine-type peptidase activity, serine hydrolase activity, aminopeptidase activity, exopeptidase activity and signalling receptor complex adapter activity. These functions are associated with enzymatic activities and signalling mechanisms (Figure3b).

**Figure 3.**
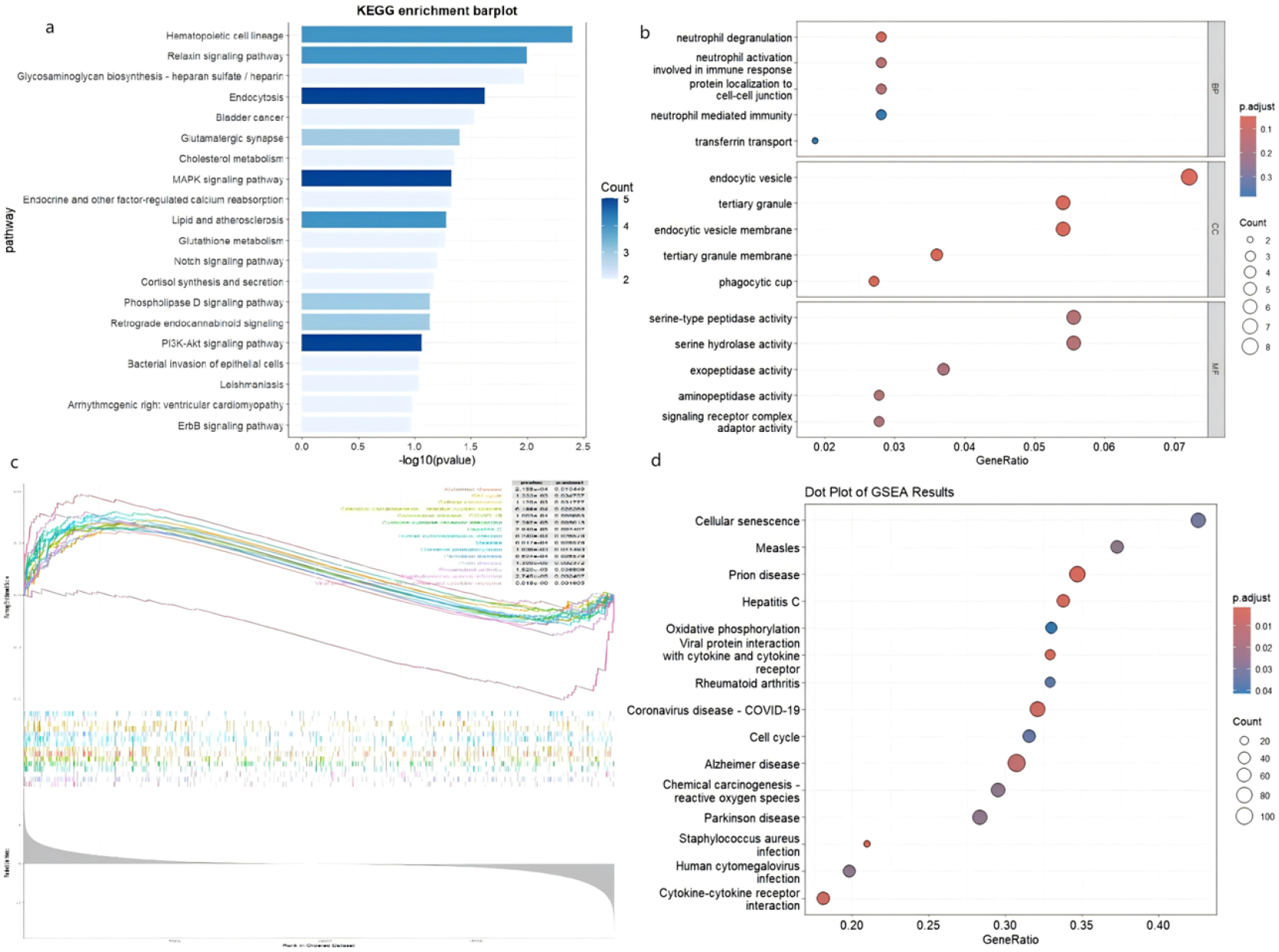
(a) KEGG Enrichment Barplot showing significantly enriched pathways with their corresponding -log10(p-value) and gene counts.(b) GO Enrichment Dotplot:A dot plot representing GO enrichment analysis, illustrating biological processes, cellular components, and molecular functions based on GeneRatio and adjusted p-values.(c) The GSEA plot illustrates enrichment scores and ranked genes for the most significant pathways.(d)Significantly enriched pathways from GSEA are summarized using GeneRatio and adjusted p-values.

The KEGG database, as the primary pathway resource, is widely used to explore the functions of genes in biological systems. For the purpose of simplification, only items with statistical significance were included. Haematopoietic cell lineages showed high statistical significance. Cholesterol metabolism, peptidoglycan lipid metabolism and calcium channels: the number of genes in these pathways is high, but the p-values are slightly higher, indicating that they are less statistically significant.

Figure3a shows the results of gene enrichment analysis for a range of biological pathways, where the length of each bar represents the number of significant genes in a particular pathway, and the colour of the bar reflects the statistical significance (p-value) of the gene enrichment. The “Haematopoietic cell lineage”, for example, shows the highest number of significant genes and the lowest p-value, suggesting that this pathway may play an important role in the biological conditions under study, especially in diseases or physiological processes involving the development of blood cells. Other pathways such as “cholesterol metabolism”, “relaxin signalling pathway”, and “calcium channel pathway” also showed some statistical significance despite the small number of genes, which may indicate their potential role in specific diseases or biological processes. Through this analysis, researchers can identify key pathways that are active in a particular biological or disease context, providing important information for further research and potential therapeutic strategies.

### GSEA Analysis

Meanwhile, GSEA (Version 4.3.2) was employed to identify key biological pathways significantly enriched in VM. This analysis focused on Alzheimer’s disease (hsa05010), the cell cycle (hsa04110), and cytokine-cytokine receptor interaction (hsa04060), as shown in Figure3c. GSEA assesses whether predefined gene sets show statistically significant, consistent differences in expression between two biological states by calculating a running enrichment score and performing a permutation test to evaluate significance. The central role of these pathways in neurodegenerative changes, cell cycle regulation, and cytokine-cytokine receptor interaction may have profound implications for the pathogenesis of Vestibular Migraine. For example, neurotransmitter changes and cell death processes involved in the Alzheimer’s disease pathway may exacerbate neurological sensitisation, whereas inflammatory modulatory effects of cytokines may play a key role in Vestibular Migraine by directly affecting pain perception and neural sensitivity. Taken together, the interactions of these pathways provide a complex molecular network perspective that contributes to a deeper understanding of the biological basis of VM and provides potential targets for the development of new therapeutic strategies.

### Identification of hub genes via machine learning

Two machine learning algorithms were applied to further filter candidate genes for vestibular migraine diagnosis (Figure 4). LASSO, a regression technique for variable selection and regularization, improves both the predictive accuracy and the comprehensibility of a statistical model [12]. RF is a powerful method that can handle both classification and regression tasks, offering high accuracy, sensitivity, and specificity without strict assumptions on variable conditions. The “glmnet” and “randomForest” R packages were used to perform LASSO regression and RF analysis. The intersection of significant genes identified by both LASSO and RF models were considered as candidate hub genes for vestibular migraine diagnosis.

**Figure 4.**
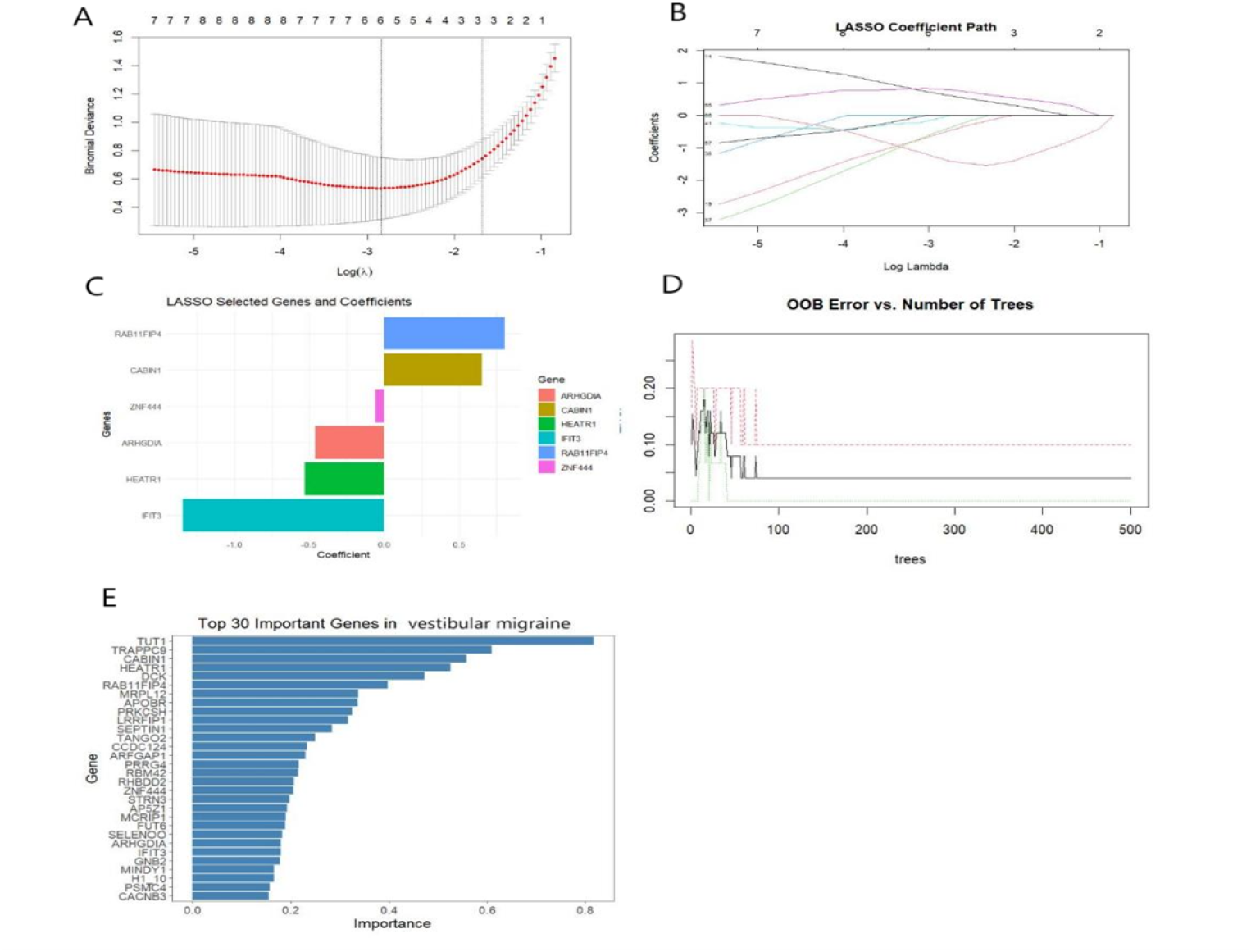
The machine algorithms for Vestibular Migraine. (A)The LASSO plot depicted the change in deviance as the log - value of the penalty parameter (lambda) varied.(B) The LASSO coefficient path plot, where the error bars represented standard errors. (C) LASSO Selected Genes and Coefficients.(D) The plot illustrated the relationship between the out-of-bag (OOB) error and the number of trees in the Random Forest model for model optimization. (E)Top 30 Important Genes in Vestibular Migraine.

In our analysis, we identified CABIN1, IFIT3, ZNF444, RAB11FIP4, ARHGDIA, and HEATR1 as key genes significantly associated with vestibular migraine. We employed both LASSO and random forest algorithms to balance prediction accuracy and computational efficiency. For the LASSO model, cross-validation identified the optimal regularization parameter λ (Figure 4A), and the coefficient paths for significant genes were plotted as λ varied (Figure 4B). In the random forest model, the optimal performance was achieved with 500 decision trees (Figure 4D). We then visualized the contribution of individual genes to the model’s accuracy, selecting the top six genes with the highest mean decrease in accuracy values for further investigation (Figure 4C, E).

Finally, the key genes identified by both LASSO and random forest models, revealing overlapping genes consistently highlighted by both methods.

Figure5a shows the differences in gene expression levels between the normal group and the vestibular migraine group. Specifically, the expression of CABIN1 and ARHGDIA was significantly higher in the normal group than in the VM group, suggesting that these genes may be downregulated in VM. On the contrary, the expression levels of IFI3 and HEATR1 in the VM group were significantly higher than those in the normal group, which may play an important role in the mechanism of VM occurrence. The expression of RAB11FIP4 showed little difference between the two groups, but was slightly higher in the VM group. The expression of ZNF444 in the normal group was significantly higher than that in the VM group, indicating its inhibitory effect in VM. The differences in gene expression patterns provide clues for understanding the potential molecular mechanisms of vestibular migraine. And the hub genes were analyzed by ROC curves in order to evaluate the diagnostic efficacy of the core genes. Among them, CABIN1 had the highest diagnostic predictive efficacy, with an AUC of 0.993(Figure5b).

**Figure 5:**
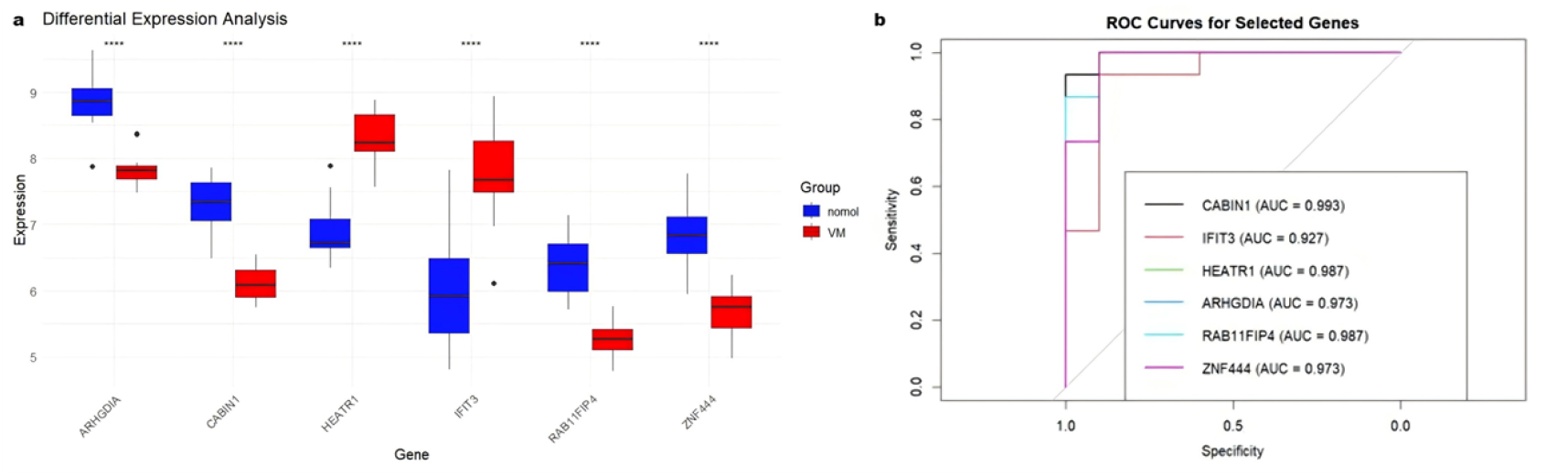
(a)This boxplot compares gene expression levels between two groups: “normal” (blue) and “VM” (red), referring to a normal group and individuals with Vestibular Migraine .(b)Diagnostic value of hub genes.

### Immune Cell Characteristics and Interrelationships in Vestibular Migraine Based on CIBERSORT Analysis

Based on the CIBERSORT algorithm for analyzing the composition of immune cells, the box plot reveals significant differences in the proportion of certain immune cell subgroups between the VM group and the normal group (Figure 6A). Specifically, the proportion of plasma cells and M0 macrophages was significantly higher in the VM group compared to the normal group, suggesting that these cells may be involved in the pathological processes of VM. Furthermore, while some T cell subsets, such as CD8 T cells, did not exhibit significant differences between the two groups, the proportion of activated CD4 memory T cells was notably elevated in the VM group (p < 0.01).

**Figure 6:**
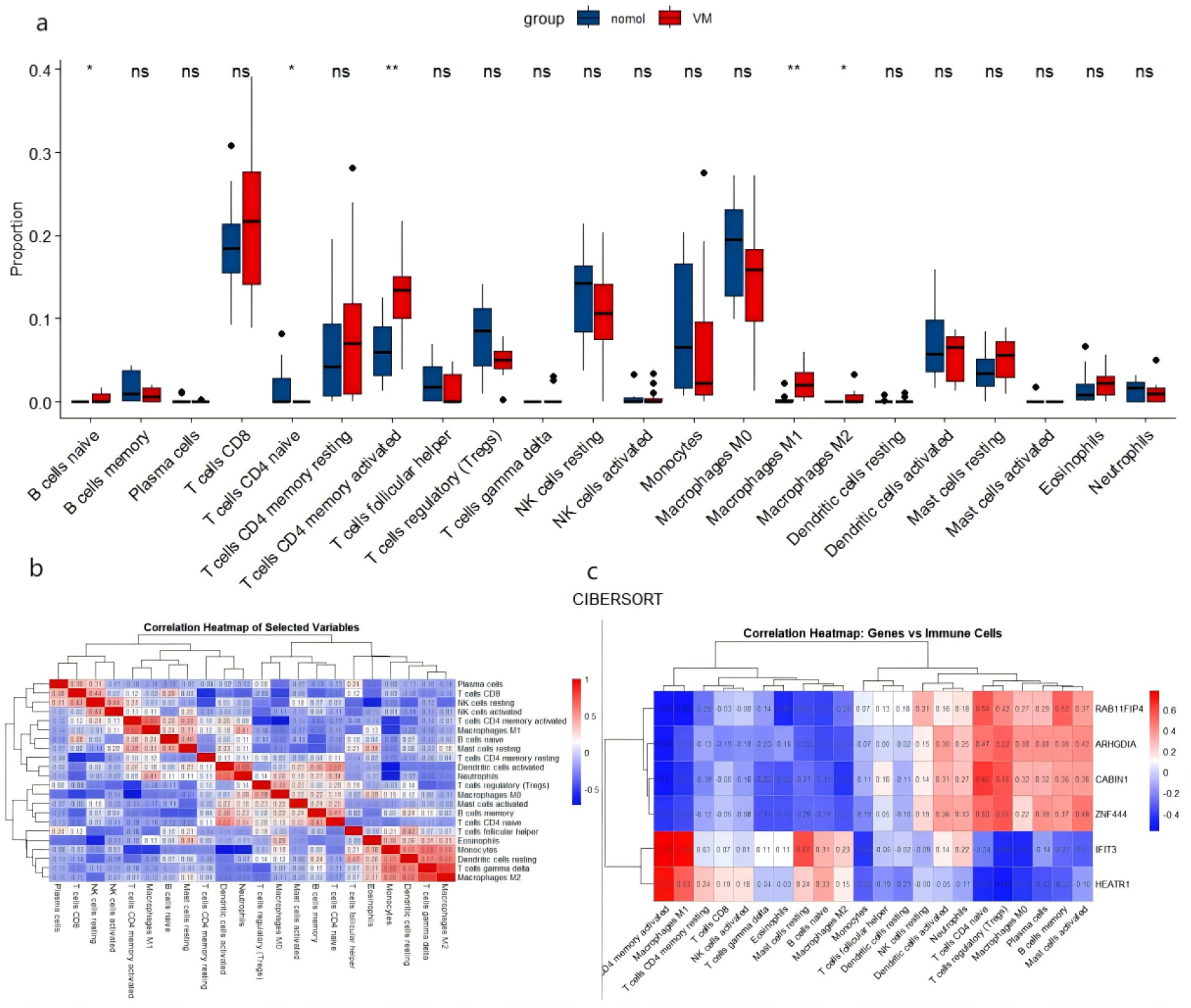
(A)This graph shows the difference in the proportion of different immune cell types between the vestibular migraine (VM, red) group and the normal group (blue) through CIBERSORT analysis. (B)This correlation heatmap shows the correlation between various immune cell subpopulations, with values ranging from -1 to 1. Red represents positive correlation, blue represents negative correlation, and the depth of the color reflects the strength of the correlation.(C)The correlation between characteristic genes and different immune cells.

Additionally, regulatory T cells showed a slight but non-significant increase in the VM group. These findings indicate that alterations in specific immune cell populations may contribute to the immune dysregulation observed in VM.

Subsequently, the interrelationships between various immune cell subpopulations were further demonstrated through correlation heatmaps(Figure 6B). This figure reveals a strong positive correlation between T cell subsets, indicating that they may jointly participate in immune regulation. In addition, M0 macrophages are negatively correlated with most other immune cell subpopulations, such as activated NK cells and T cells, suggesting that they may participate in regulating inflammatory responses through different immune pathways. Figure 6C further illustrates the correlation between characteristic genes (such as CABIN1, ZNF444) and various immune cell infiltration ratios, with CABIN1 showing a strong positive correlation with cells such as Macrophages M0 and Neutrophils, suggesting that these genes may play a key role in regulating immune cell infiltration.These findings provide a new perspective for understanding the immune mechanisms of VM.

### TF-gene interaction

Figure 7 shows the interaction network between genes. The red nodes represent key genes, which have more connections with other genes in the network. The resulting TF - gene interaction network comprised 36 nodes and 63 edges.

**Figure 7:**
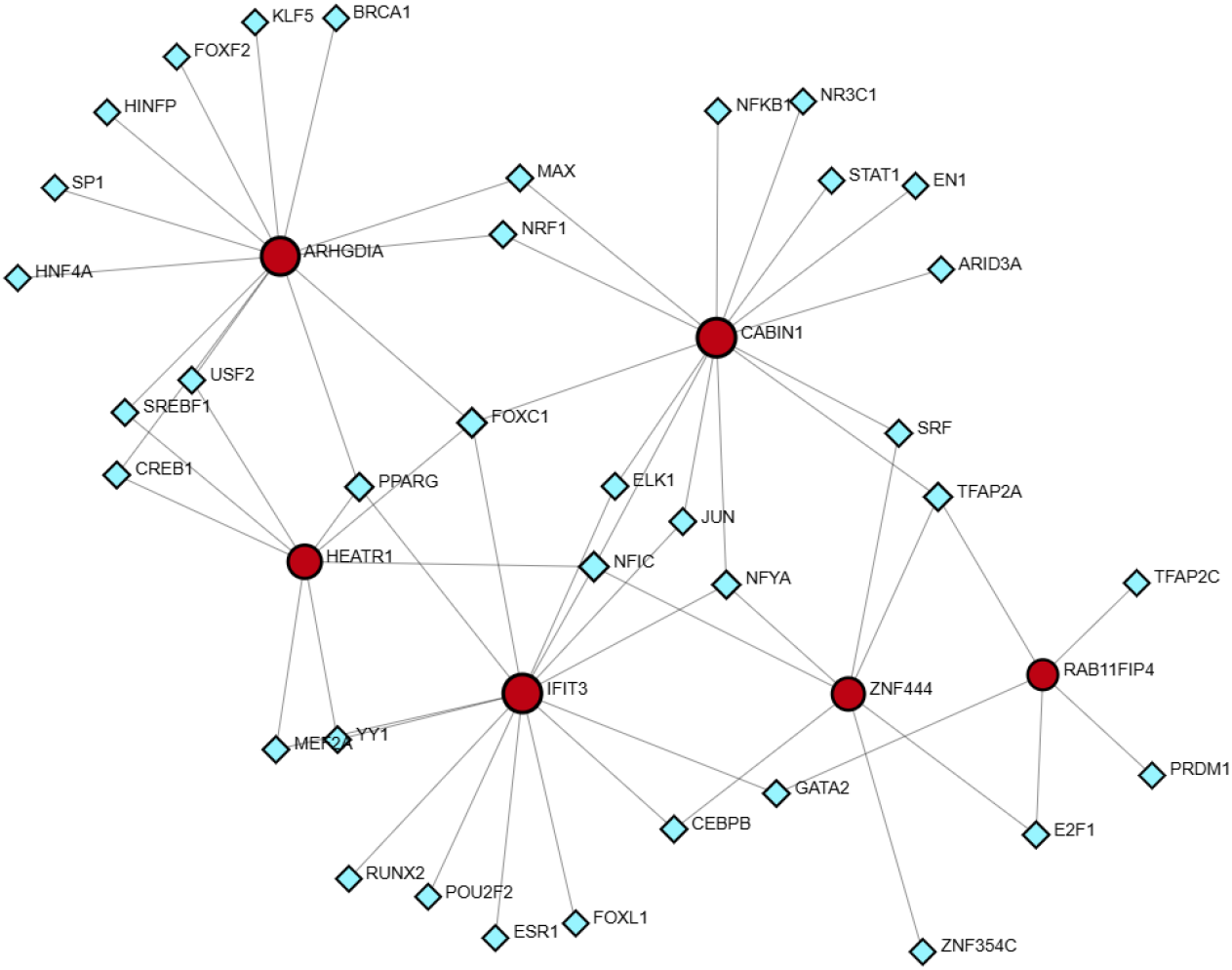
TF - gene interaction network featuring 6 hub genes. Red circles signify the 6 hub genes, while blue diamonds denote transcription factor genes that are connected to the hub genes.

FOXC1 interacted with four hub genes: ARHGDIA, CABIN1, HEATR1, and IFIT3. NFYA engaged with three hub genes: CABIN1, IFIT3, and ZNF444. TFAP2A interacted with three hub genes: CABIN1, ZNF444, and RAB11FIP4. Notably, the three transcription factors had CABIN1 as a common hub gene.

### Molecular docking simulation

Several drugs, such as Acetaminophen, bisphenol A, and Phenylephrine, were identified through the CTD.The potential therapeutic mechanisms of these drugs were explored through molecular docking simulations. CABIN1 protein was docked with the three compounds respectively to evaluate their binding affinities as potential therapeutic targets.Figure 8A shows a schematic diagram presenting the interaction pattern between the expression product of the CABIN1 gene and Acetaminophen. It clearly indicates the interaction distances (3.1 Å and 2.9 Å) between the key amino - acid residues (such as TYR - 1264 and ALA - 1260) and the ligand. Figure 8B is a three - dimensional structural schematic diagram presenting the interaction pattern between the gene and bisphenol A. The light - green protein exhibits a spatial conformation of helices and coils. The distances (3.4 Å, 3.1 Å, 2.5 Å) between amino - acid residues such as “LYS - 1162”, “MET - 1761”, and “ASP - 1762” and the ligand are also shown. Figure 8C demonstrates the interaction between CABIN1 and Phenylephrine. The light - green protein shows a complex morphology of helices and coils. Multiple amino - acid residues labeled in dark pink, such as “TYR - 1244” and “PHE - 1299”, and the corresponding distance values (such as “2.6”, “3.3”, etc., in the unit of Å) with the ligand are presented. These distance information are of great value for elucidating the interaction mechanism between the two and the stability of the complex, providing a structural reference for relevant drug research and development and other studies.

**Figure 8:**
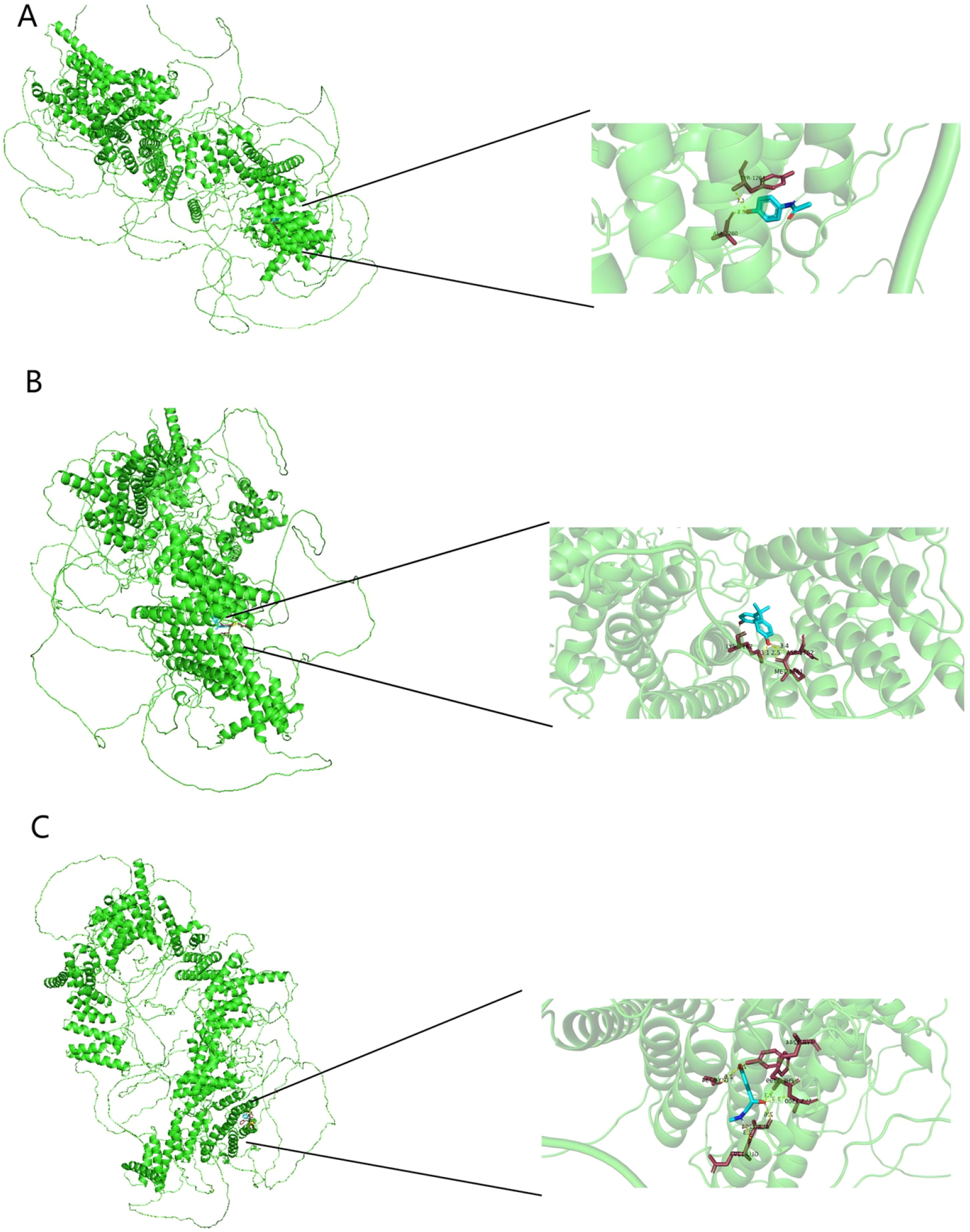
(A)Schematic of Molecular Docking Interaction Pattern and Key - site Distances between CABIN1 and Acetaminophen;(B)Three - dimensional Structure of the Interaction between CABIN1 and Bisphenol A and Display of Amino - acid – ligand Distances;(C)Binding Pattern of CABIN1 and Phenylephrine and Presentation of Key Amino - acid Residue Distances”

## Discussion

Many studies have proposed that vestibular migraine is an ion channel gene defect disease, and the discovery of effective therapeutic targets for Vestibular Migraine is crucial due to the lack of effective treatments[13, 14, 15]. However, with the continuous research on Vestibular Migraine over the last decade and updated bioinformatic analysis methods, it is hoped that this study will provide new insights and hope for the treatment of the disease, offering valuable clues or methods for the diagnosis or treatment of Vestibular Migraine.

Increasing evidence highlights the pivotal role of inflammation in the pathogenesis of Vestibular Migraine[16]. Inflammatory processes, characterized by glial cell activation and leukocyte infiltration, are believed to contribute significantly to Vestibular Migraine’s development and maintenance[17]. Pro-inflammatory cytokines and chemokines are central to these processes, affecting neural pathways involved in pain perception and vestibular function. Inflammatory mediators can lead to sensitization of neural pathways, contributing to the chronicity and severity of VM symptoms. Additionally, inflammation is linked to other common manifestations in Vestibular Migraine patients, such as mood disorders and cognitive dysfunction, further complicating the clinical picture[18]. Our study identified KEGG involved in these inflammatory processes, suggesting potential diagnostic and therapeutic targets.

In this study, we searched the GEO dataset and found only one expression profile of GSE109558 associated with human gene-based Vestibular Migraine. GO functional and KEGG enrichment analyses showed that most of the overlapping degs were mainly enriched in the immune-inflammatory response and the calcium channel pathway, inducing both peripheral and central sensitisation. Six important genes (“IFIT3”, “RAB11FIP4”, “ZNF444”, “ARHGDIA”, “HEATR1”, “CABIN1”) were identified by lasso regression and random forests, with “CABIN1” being the most important gene. CABIN1 (Calcineurin-binding protein 1) plays several key roles in the nervous system, especially in neural development and neuronal function regulation. It has been found that Cabin1 is an essential gene for neural development in mice and is enriched in the CNS region of developing zebrafish[19]. The researchers determined the spatiotemporal expression of Cabin1 mRNA during CNS development, particularly in the later stages of nervous system development[20]. In certain regions of the CNS such as the olfactory system and the cerebellum, Cabin1 expression overlapped with regions of expression of proteins known to interact with Cabin1 (MEF2 and/or calmodulin phosphatase). In addition it has been shown that reduced expression of Cabin1 protein leads to diminished auditory and vestibular function in developing zebrafish.

Cabin1 is a repressor of MEF2- (myocyte enhancer factor 2) and calmodulin phosphatase-mediated transcription in the immune system. Calmodulin phosphatase is a calcium/calmodulin-dependent serine/threonine phosphatase involved in a variety of cellular functions, including synaptic transmission and plasticity in neurons[21].CABIN1 regulates neuronal signalling and survival by binding to and inhibiting the activity of calmodulin phosphatase.Cabin1 is cleaved by calmodulin during apoptosis, resulting in the loss of CABIN1 inhibition of calmodulin phosphatase inhibition and activation of calmodulin phosphatase, which in turn mediates calcium-triggered cell death. This mechanism is particularly important in neurodegeneration caused by neuronal excitotoxicity. Hyperactivation of calmodulin phosphatase may lead to greater sensitivity of neurons to excitatory amino acids (e.g., glutamate), increasing excitotoxicity, which in turn triggers neuronal injury and death.Aberrant expression of CABIN1 may be associated with a variety of neurological disorders. Overactivation of calmodulin phosphatase is associated with neurodegenerative diseases (e.g., Alzheimer’s disease, Parkinson’s disease) and neuronal excitotoxicity injury.Decrease in the expression level of CABIN1, an inhibitor of calmodulin phosphatase, may lead to the overactivation of calmodulin phosphatase, which may trigger or exacerbate these diseases.

Taken together, CABIN1 acts in the nervous system mainly by inhibiting the transcriptional activities of calmodulin phosphatase and MEF2. CABIN1 plays a key regulatory role in neuronal development, synapse formation, and neuronal survival, and its low expression may lead to increased excitotoxicity of neurons, aberrant synaptic plasticity, and an increased risk of neurological disorders[22]. Our mechanistic learning allowed us to find that CABIN1 is low expressed in patients with VM, thus explaining the abnormalities of the CABIN1 gene in patients with vestibular migraine, leading to overactivation of calmodulin phosphatase, altered Ca2+ concentrations, oxidative stress and inflammatory mediators mediating disease. Calcium channel blockers (flunarizine, etc.)[23] have a role in the treatment of VM, and future studies should further explore the specific mechanisms of CABIN1 in the nervous system with the aim of providing new targets for the treatment of related diseases.

This study explores the molecular mechanisms underlying Vestibular Migraine (VM) and identifies key feature genes as potential diagnostic biomarkers. Using gene expression data (GSE109558) and various bioinformatics methods (such as limma, WGCNA, GO analysis, KEGG pathway analysis, and GSEA), the study found that these differentially expressed genes are associated with biological processes like inflammation and calcium ion channel regulation. Further feature selection was carried out using lasso regression and random forest methods. Immune cell infiltration analysis was performed using CIBERSORT, and Spearman’s correlation analysis was used to explore the relationships between differentially expressed genes and immune cells. Molecular docking simulations were employed to explore their potential therapeutic mechanisms.While the findings lay a solid foundation for future research, further experimental validation and clinical studies are needed to confirm the diagnostic and therapeutic potential of these biomarkers and to optimize treatment approaches for VM patients.

## Conclusion

In summary, after bioinformatics analysis and selection of machine learning methods, CABIN1 has been identified as a potential key gene and potential diagnostic biomarker for VM. To our knowledge, research on human CABIN1 is limited. This study identified CABIN1 as a potential key gene for VM , which may provide a new target for the diagnosis and treatment of VM. In the future, studies with larger sample sizes should be conducted to further validate our findings, particularly to confirm the robustness of the identified biomarkers and therapeutic targets, as well as to evaluate their clinical applicability and efficacy across different populations.

## Abbreviations

VM: Vestibular Migraine
CABIN1: Calcineurin-binding protein 1
DNA: Deoxyribonucleic Acid
RNA: Ribonucleic Acid
DEGs: Differentially expressed genes GO:Gene ontology
KEGG: Kyoto encyclopedia of genes and genome
CNS: Central Nervous System
SNP: Single Nucleotide Polymorphism
IFIT3: interferon induced protein with tetratricopeptide repeats 3
HEATR1: HEAT repeat containing 1
ARHGDIA: Rho GDP dissociation inhibitor alpha
RAB11FIP4: RAB11 family interacting protein 4
ZNF444: zinc finger protein 444

## Author contributions

YXT designed the study. YXT and LZJ performed data analysis. LYL drafted and revised the manuscript. YXT and LZJ contributed equally to this work. All authors (YXT, LZJ, and LYL) read and approved the final manuscript.

## Funding Declaration

This work was partially supported by research funding provided by The Third Affiliated Hospital of Guangzhou Medical University, with additional costs covered by the authors.

## Ethics Declaration

All data used in this study were obtained from public databases; therefore, this study did not include any intervention experiments related to animals or humans.

## Acknowledgments

The authors thank GEO for providing data.

## Data availability statement

The data that support the findings of this study are available in The National Center for Biotechnology Information at https://www.ncbi.nlm.nih.gov/sites/GDSbrowser/, reference number GSE109558.

## Clinical trial number

Not applicable. This study is based on analysis of data from a public database and does not involve new clinical trials or patient interventions.

## Notes

### Competing Interest Statement

The authors have declared no competing interest.

